# A digital approach to protein identification and quantity estimation using tandem nanopores, peptidases, and database search

**DOI:** 10.1101/024158

**Authors:** G. Sampath

## Abstract

A digital approach to protein identification and quantity estimation using electrical measurements and database search is proposed. It is based on an electrolytic cell with two (three) nanopores and one (two) peptidase(s) covalently attached to the *trans* side of a pore. An unknown protein is digested by a reagent or peptidase into peptides ending in a known amino acid; the peptides enter the cell, pass through the first pore, and are fragmented by a high-specificity endopeptidase. The second enzyme, if present, is an exopeptidase that cleaves the fragments into single residues after the second pore. Level transitions in an ionic blockade or transverse current pulse due to residues in a fragment or individual pulses due to single residues are counted. This yields the positions of the endopeptidase’s target in the peptide, and, together with the peptide’s terminal residue, a partial sequence. Search through the Uniprot database for such sequences identifies over 90% of the proteins in the human proteome. The percentage can be increased by repeating the procedure with other reagents and cells specific to other residues, close to 100% may be possible. Sample purification to homogeneity is not required as the method applies to an arbitrary mixture of proteins; the quantity of a protein in the sample is estimated from the number of identifying peptides sensed over a long run. A Fokker-Planck model gives minimum enzyme turnover intervals required for ordered sensing of peptide fragments. With thick (80-100 nm) pores, required pulse resolution times are within the capability of CMOS detectors. The method can be implemented with existing technology; several related issues are discussed.

## 1 Introduction

Currently sequencing and identification of proteins are largely based on the established techniques of Edman degradation,^1,2^ gel electrophoresis,^1,2^ and mass spectrometry.^3^ Sequencing tends to use bulky or expensive devices and/or time-consuming procedures; this has led to efforts aimed at developing portable/hand-held low-cost fast-turnaround devices.^4^,^5^ In particular, nanopores have been investigated for their potential use in the analysis and study of DNA^6^ and proteins/peptides.^7^-^10^ Recently a tandem electrolytic cell with cleaving enzymes was proposed for sequencing of DNA^11^ or peptides.^12^ It has two single cells in tandem, with the structure [*cis*1, upstream pore (UNP), *tra*ns1/*cis*2, downstream pore (DNP), *trans*2]. An enzyme covalently attached to the downstream side of UNP successively cleaves the leading monomer in a polymer threading through UNP; the monomer translocates through DNP where the ionic current blockade it causes is used (along with other discriminators^12^) to identify it. With DNA the enzyme is an exonuclease,^11^ with peptides it is an amino or carboxy peptidase.^12^ The process is label-free and does not require immobilization of the analyte.

Here a low-cost alternative for protein identification is proposed in which a partial sequence is obtained for a peptide and used to identify the protein by comparison with sequences in a target proteome. Partial sequencing is done with a tandem cell^11^,^12^ and an endopeptidase (*Method 1*), or a double tandem cell, endopeptidase, and exopeptidase (*Method 2*). The first enzyme breaks the peptide into fragments, the second breaks fragments into residues. The fragments/residues translocate through a pore and cause ionic current blockades or modulate a transverse current across the pore;^6^ the pore/transverse current pulses or level transitions within are counted. With an endopeptidase specific to an amino acid a list of integers corresponding to the positions of the amino acid in the peptide sequence is produced. Along with the peptide’s terminating residue this yields a partial sequence, which is compared with sequences in a protein database such as Uniprot. Calculations show that with this approach at least 93% of the proteins in the human proteome can be identified. Purification of the assay sample to homogeneity is not required as the method applies equally to a mixture of proteins, with the quantity of a protein in a mixture being estimated from the number of its identifying peptides. The proposed approach may be extended to include modified amino acids.

This is a digital technique based on pulse counting, it differs from other nanopore sequencing and identification techniques based on analog measurements of pulse magnitude or width (equivalently analyte residence time in a nanopore) of a pore ionic or transverse current.^6^,^11^,^12^ The approach is similar in some ways to a recent theoretical proposal^13^ for peptide identification that combines fluorescent labeling with a series of Edman degradation cycles to identify the N-terminal residues of an immobilized peptide, followed by database search. In contrast, the proposal presented here does not use analyte immobilization, labeling, or repeated wash cycles.

## 2 Protein identification and quantity estimation: method and materials

An unknown protein P is identified in five stages:

1. Fragment P into peptides ending in amino acid X_0_.
2. Break peptide into fragments ending in amino acid X_1_ ≠ X_0_ (*Methods 1* and *2*). Break a fragment into individual residues (*Method 2*).
3. Find number of residues in each fragment obtained from Stage 2.
4. Assemble partial sequence of peptide; mark unknown residues with wild card *.
5. Match partial sequence with sequences in proteome of interest and identify P, hopefully uniquely.

**Stage 1**. A highly specific chemical or peptidase targets a single amino acid X_0_. Examples include cyanogen bromide, which cleaves after methionine (M), and GluC protease, which cleaves after glutamic acid (E). (For a list of selected reagents/peptidases see Table A-4 in the Appendix, which has been adapted from an online review^14^ that includes a list of comprehensive references.) This results in the protein being broken into peptides that end in X_0_.

**Stage 2**. The positions of occurrence of a single residue X_1_ in a peptide are obtained by targeting it with a highly specific peptidase in a tandem cell. Peptidases with high specificity include GluC (which targets E), ArgC proteinase (arginine, R), AspN endopeptidase (aspartic acid, D), and LysC lysyl endopeptidase (lysine, K). See Table A-4 in the Appendix. In turn a peptide fragment can be cleaved into a series of individual residues by an exopeptidase capable of cleaving a wide range of residue types at the carboxyl or amino end. Examples include Carboxypeptidase I (CPD-Y), Carboxypeptidase II (CPD-M-II), and Leucine Aminopeptidase (LAP).^15^-^17^

**Stage 3**. A tandem cell is used to count level transitions in a pore ionic current or transverse current pulse that is modulated by a fragment translocating through a nanopore, or individual such pulses due to single residues cleaved from the fragment. Two methods are considered.

**Figure 1.**
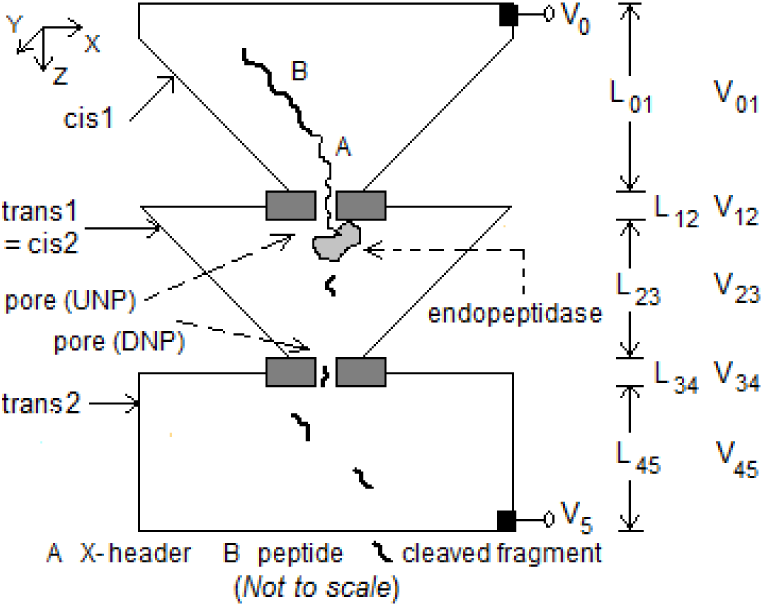
Tandem cell for peptide sequencing. Dimensions: 1) *cis*1: box of height 1 μm tapering to 100 nm^2^; 2) membrane with UNP: length 10-20 nm, diameter 10 nm; 3) *trans*1/*cis*2: box of height 1 μm tapering from 1 μm^2^ to 10 nm^2^; 4) membrane with DNP: length 10-20 nm, diameter 3 nm; 5) *trans*2: box of height 1 μm, side 1 μm. Endopeptidase covalently attached to downstream side of UNP. Electrodes at top of *cis*1 and bottom of *trans*2. V_05_ = ∼115 mV.

**Figure 2.**
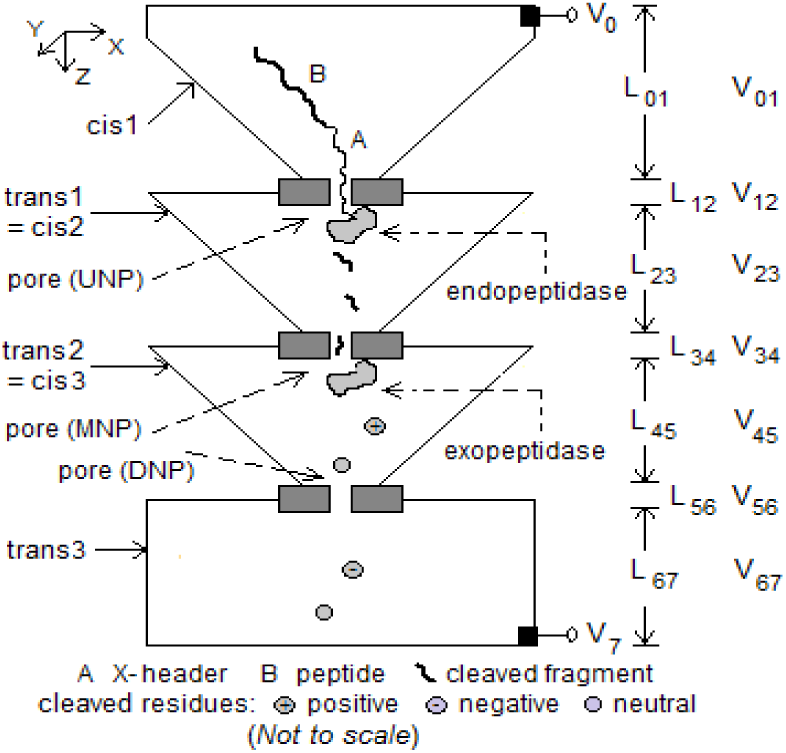
Double tandem cell for peptide sequencing. Dimensions: 1) *cis*1: box of height 1 μm tapering to 100 nm^2^; 2) membrane with UNP: length 10-20 nm, diameter 10 nm; 3) *trans*1/*cis*2: box of height 1 μm tapering from 1 μm^2^ to 10 nm^2^; 4) membrane with MNP (middle nanopore): length 10-20 nm, diameter 3 nm; 5) *trans*2/*cis*3: box height 1 μm tapering from 1 μm^2^ to 10 nm^2^; 6) membrane with DNP: length 40-50 nm, diameter 3 nm; 7) *trans*3: box of height 1 μm, cross-section 1 μm^2^. Endopeptidase (exopeptidase) covalently attached to downstream side of UNP (MNP). Electrodes at top of *cis*1 and bottom of *trans*3. V_07_ = ∼180 mV.

In *Method 1* the structure in Figure 1 is used. A peptide with a poly-X header (X = one of the charged amino acids: Arg, Lys, Glu, Asp; the charge on X depends on the pH value) is drawn into UNP by the electric field due to V_05_ (= ∼110 mV), most of which (∼98%) drops across the two pores.^6^ An endopeptidase specific to amino acid X_1_ attached downstream of UNP cleaves the peptide after (or before) all n (≥ 0) points where X_1_ occurs. The resulting n+1 fragments (the first of which ends in X_0_) translocate to and through DNP, where level crossings in the resulting pore ionic current blockade or a transverse current across DNP are used to count the residues in a fragment.

In *Method 2* the double tandem cell in Figure 2 is used. A peptide is cleaved into fragments by an endopeptidase as in *Method 1*. An exopeptidase (amino or carboxy) covalently attached downstream of the middle nanopore (MNP) cleaves successive residues in a fragment, the residues translocate through DNP and blockade the pore ionic current or modulate the transverse current. The resulting single pulses are counted.

In both methods a tandem cell specific to amino acid X_1_ produces an ordered list of integers equal to the lengths of successive fragments in which the last residue is the target X_1_ (except the first fragment, which ends in X_0_). If the cell generates a single integer, the target is not in the peptide.

**Stage 4**. A partial sequence S_X_’ for peptide P_X_ is partially assembled as follows:

- *Step 1*: Replace fragment lengths from cell with cumulative lengths (= target positions of X_1_ in the peptide).
- *Step 2*: Invert position-identity pairs to obtain S_X_^’^ (with X_0_ in the last position).
- *Step 3*: Insert wild card * in all unknown positions in S_X_^’^.

The resulting sequence is a partial sequence containing X_1_ and ending in X_0_.

**Stage 5**. Standard string matching algorithms can be used to search for S_X_^’^ among sequences in a protein database such as Uniprot^18^ or PDB. More general matching algorithms^19^ may be used for error detection and correction.

**Cell output**. The output for a given X_0_-X_1_ pair is a series of partial sequences containing X_1_ and ending in X_0_ corresponding to peptides from the unknown protein entering the cell in some random order. If the assay sample consists of a mixture of proteins, the output corresponds to peptides from any of the proteins in the mixture in any order. As discussed below, by repeating the procedure with different X_0_ and X_1_, different partial sequences can be obtained for the same protein and used to increase the identification rate.

### 2.1 Database search and results

A partial sequence obtained in the procedure described above can be used to identify the container protein based on precomputing all identifying peptides for each protein in a given proteome. Identification is then a matter of matching the partial sequence output by a cell with a precomputed sequence. The process is illustrated with the human proteome (Uniprot database id UP000005640, manually reviewed subset with 20207 sequences) for the following set of cleavage targets: X_0_ = M (first stage), and X_1_ = R (second stage).

#### Precomputation

1. All subsequences of proteins in the protein that end in M are extracted, they correspond to the peptides generated by the action of cyanogen bromide (see above). Every one of these peptides has exactly one M which is also the last residue in the peptide.
2. Enter wild card * into every position in every peptide sequence in the database where R and M do not occur.
3. Each subsequence (peptide) is compared with every other reduced subsequence from Step 2. If there is no match mark the peptide as a unique identifier.

The resulting data are available for download; see Appendix.

#### Identification

1. Read output data from Stage 4 above, this corresponds to a partial sequence for a peptide.
2. Compare partial sequence with sequences of identifiable proteins in precomputed database; output identity of (unique) container protein if there is a match.

The percentage of proteins with at least one identifying subsequence (created by cyanogen bromide in Stage 1 and cleaved after each occurring R by ArgC in Stage 2) is found to be 90.49%. It may be increased by using other combinations of cleaving chemical/enzyme and peptidases (Table A-4, Appendix). Figure 3 shows the distribution of the number of proteins vs the number of identifying peptides in a protein for two sets of cleavage choices in Stages 1 and 2. The total coverage is the union of the sets of proteins with at least one unique identifier obtained from all X_0_-X_1_ pairs used.

**Figure 3.**
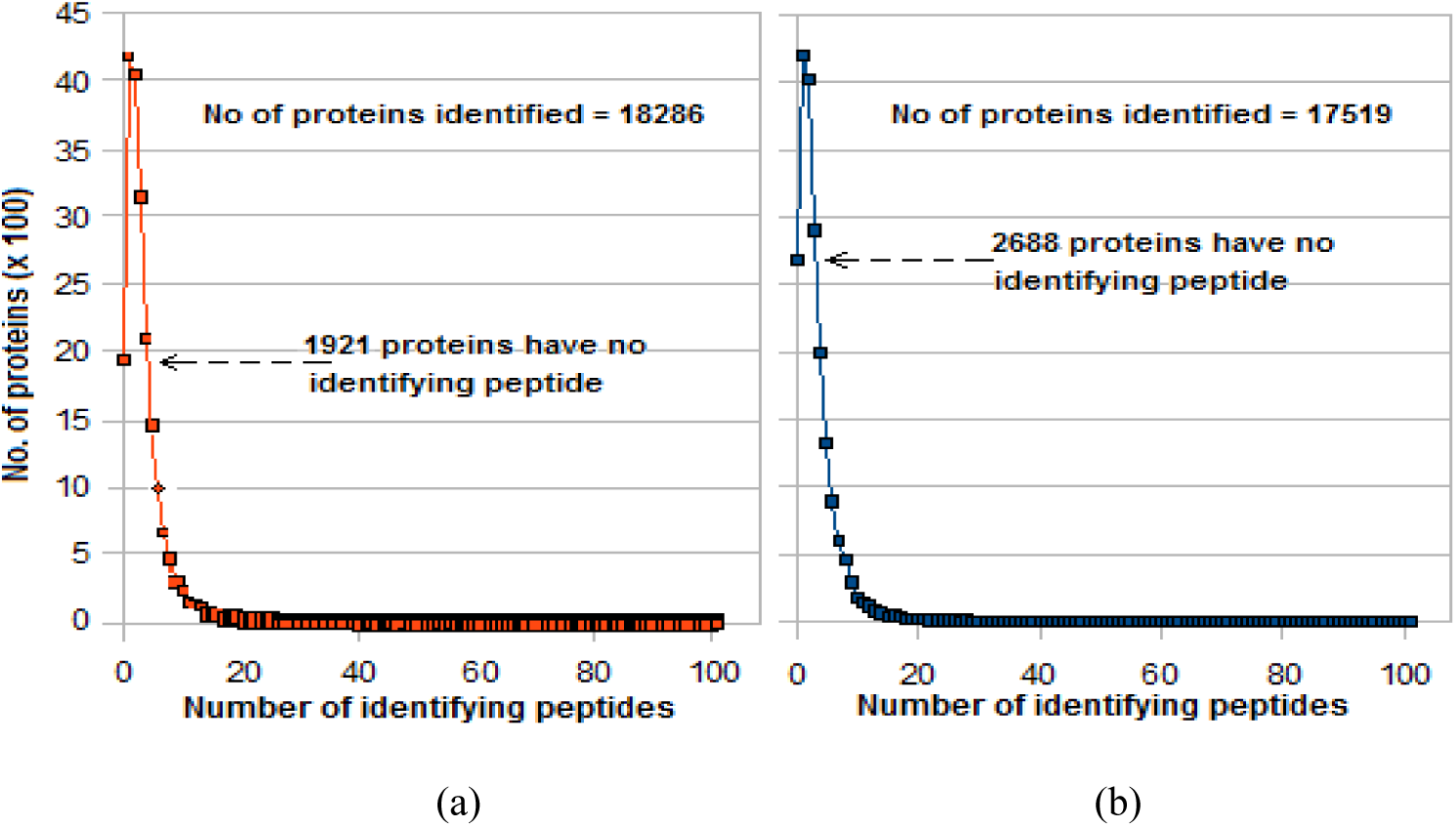
Number of proteins with a given number of identifying peptides in the human proteome (UP000005640, manually reviewed subset with 20207 sequences): (a) cleave M in Stage 1 and R in Stage 2; (b) cleave Y in Stage 1 and R in Stage 2

Table 1 shows the increase when sequencing is done twice by targeting M or Y in Stage 1, with R targeted both times in Stage 2.

**Table 1:** **Increase in number of identifiable proteins from multiple runs with different cleavage targets (Total number of proteins in human proteome** UP000005640 **= 20207)**

| Run no. | Target in Stage 1 | No of peptides created in Stage 1 | Target in Stage 2 | No. of proteins identified / not identified | Not identified in this run, identified in the other | Total proteins identified in both runs / not identified |
| --- | --- | --- | --- | --- | --- | --- |
| 1 | M | 261048 | R | 18286 = 90.49%<br>1921 = 9.51% | 522 | 18808 = 93.07%<br>1399 = 6.93% |
| 2 | Y | 321376 | R | 17519 = 86.69%<br>2688 = 13.30% | 1288 |  |

### 2.3 Estimating the quantity of a protein in a mixture

Consider an assay of a mixture of proteins given by {(N_i_, P_i_, I_i_): i = 1, 2,…} where N_i_ is the number of molecules of the i-th protein in the mixture, P_i_ the number of peptides per molecule of the protein (this is equal to the number of peptides created in Stage 1 from a single molecule), and I_i_ (0 ≤ I_i_ ≤ P_i_) the number of identifying peptides per molecule. For a given chemical agent/peptidase in Stage 1 the P_i_s are known, and for the set of peptidases used in the cells in Stage 2 the I_i_s are known by computation. N_i_ is the desired unknown. For example, with M targeted in Stage 1 and R in Stage 2, for the protein P31946 in the human proteome P_i_ = 9 and I_i_ = 1. Peptides generated in Stage 1 from the mixture enter a cell in some random order and are partially sequenced and used to identify their container proteins. Let the total number of peptides processed in the run be N_total_ and the number of peptides in protein i that have been identified in the run so far I_i-measured_. If peptide entry into a cell is totally random, then after a sufficiently long run N_i_ can be estimated as

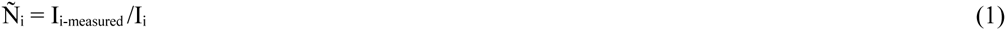

The corresponding fraction is

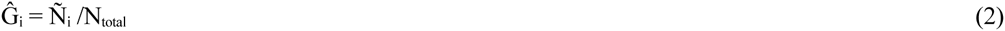

The number of peptides that do not yield identifying information is

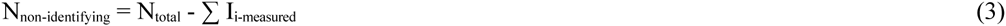

where the summation is over all the identified proteins. This number includes peptides that are found in more than one protein and may also include impurities in the assay sample. If the sample is contaminated there seems to be no easy way to separate the two so unidentified sample proteins that are not impurities remain unestimated (even though the percentage is likely small).

## 3 Necessary conditions for ordered recognition of fragments from a peptide

Nanopore-based sequencing relies on the ability to measure changes in current flow when an analyte molecule is present. This current may be an ionic current from *cis* to *trans*, a transverse electronic current across the pore membrane, or a transverse tunneling current across a gap in the membrane.^6^ The measurement ability is closely related to the bandwidth of the detector, see discussion in Section 4 below.

Since the charge carried by a peptide is highly variable and may be negative, 0, or positive, the two methods described above rely on diffusion as the primary mechanism for translocation of a fragment or residue, modified by the drift field. They are studied through the properties of the basic tandem cell, which has been modeled with a Fokker-Planck equation.^11^,^12^ Central to the model is the solution of a boundary value problem in which the *trans* side of a pore is viewed as a reflecting boundary for a cleaved fragment or residue, so the net diffusion tends to be in the *cis*-to-*trans* direction (with V_05_, V_07_ > 0). The main quantities of interest are the mean E(T) and variance σ^2^(T) of the time T taken by a particle to translocate through a *trans* compartment or pore of length L (in the latter case it is ≈ the width of the pore ionic blockade or transverse current pulse) and with applied potential difference of V. Expressions for E(T) and variance σ^2^(T) are given in Section A-1 of the Appendix, detailed derivations may be found elsewhere.^11^,^12^

For fragments to yield position information for residues targeted by the endopeptidase, they must be sensed in their natural order in the second pore in *Method 1* (or residues from fragments in the third pore in *Method 2*). The following is a set of necessary conditions for this to happen.

*Necessary conditions*

*C1*: a) At most one cleaved fragment may occupy DNP (*Method 1*) or MNP (*Method 2*) at any time; b) At most one cleaved residue may occupy DNP (*Method 2*).

*C2*: a) Cleaved fragments (*Method 1*) or residues (*Method 2*) must arrive at DNP in sequence order; b) Cleaved fragments must arrive at MNP in sequence order (*Method 2*).

*C3*: a) A residue translocating through DNP must have a blockade pulse width > T_detector_ (*Method 2*); here T_detector_ = time resolution of the detector circuit (= ∼1 μs with CMOS circuits^20^).

b) A fragment with L_f_ residues must have a blockade pulse width in DNP > L_f_T_detector_ (*Method 1*).

The following are some factors to consider in ensuring that these conditions are satisfied.

- The pore ionic blockade or transverse current pulse width, which is effectively the fragment or residue’s residence time in DNP. It is approximated by the mean translocation time through DNP in both methods.
- The charge carried by a peptide fragment (and hence its mobility μ). As it depends on the constituent amino acids it has a wide range of values, which directly affects the translocation time (see Section A-2 in the Appendix for the relevant equations). Thus fragments with high negative charge have very high speeds of translocation which may result in misses (‘deletes’), while those with high positive charge are ‘lost’ to diffusion because they are too slow. Figure A-1 in the Appendix shows the frequency distribution of all 20^7^ peptides of length L_F_ = 7 as a function of μ or μ/D at pH = 7 (physiological pH), where D is the diffusion constant of the fragment. (Note the multimodal shape and slight negative skew.) These distributions are used in the Appendix to estimate the percentage of misses (deletes) and losses.

*Satisfying the necessary conditions*

- *C1* and *C2* can be satisfied by requiring the enzymes to cleave at a given minimum rate. Enzyme reactions being stochastic processes, reaction rates are random variables with a distribution of values. The minimum rates required are estimated using standard statistical methods.
- *C3* can be satisfied through the use of a sufficiently thick pore. Thus the pore ionic blockade or transverse modulated current pulse width is proportional to the square of pore length (Equations A-1 through A-4), so a thicker pore can significantly increase translocation times and thus lower the required bandwidth (or equivalently increase the resolution time needed to sense the pulse). This is contrary to the usual practice of using thinner pores to achieve better discrimination,^6^ but is appropriate here because residues do not have to be identified, they only have to be counted. (A side benefit of this is that thick synthetic pores are usually easier to fabricate than thin ones.^21^) See Discussion in Section 4.

With suitable values for the pore length, applied voltage, and peptide length all three conditions can be satisfied with T_detector_ = ∼1 μs. See Appendix for details.

*Comparing the two methods*

*Method 1* has a more compact physical structure and uses only one enzyme, but the need to recognize transitions in a blockade pulse due to a fragment reduces the maximum length that can be determined accurately. The ionic current is also lower. *Method 2* can use a shorter (that is, thinner) DNP and a higher potential difference (leading to a higher ionic current). (This is not as serious a problem with transverse currents, which are on the order of nA,^21^ compared with at most 100s of pA with ionic currents.^6^) However, as noted earlier, the reaction time required of the endopeptidase is larger because of the need to sense more than one residue in DNP; also both the endopeptidase and the exopeptidase need to cleave at a sufficiently low rate and in synchrony. Notice that in *Method 2* even if the exopeptidase is inefficient and does not cleave after every single residue, the number of residues would be counted correctly if DNP senses transitions between residues in a pulse due to a fragment with more than one residue.

## 4 Discussion

Implementation of the proposed scheme appears both feasible and practical given that the required chemistry, nanopore fabrication technology, detector electronics (with needed bandwidth), and database search methods are all currently available. Some related issues are considered next.

1) *Counting pulses or transitions in a pulse*. On the face of it counting transitions in a pulse due to residues in a fragment would appear to be easier with the following methods: a) using single-atom thick graphene^22^ or molybdenum disulphide (MoS_2_) sheets,^23^ both of which make counting of transitions easier; b) detecting level crossings in a transverse electronic or tunneling current pulse across graphene or silicon gaps;^24,25^ and c) using a narrow biological nanopore like MspA, which has a constriction in its short stem^26^ that may aid in recognizing the transitions. However all of these methods would require bandwidths in the tens of MHz if directly used in the approach described here. To bring the bandwidth down to 1-2 MHz (corresponding to a pulse width resolution of ∼1 μs), thick pores may be considered, as discussed in Section 3. With silicon compounds like Si_3_N_4_ thick pores (50-100 nm) are actually easier to manufacture than thin ones.^21^,^27^ With graphene, hourglass-shaped pores may be fabricated from graphite (which is a stack of graphene layers^28^), but stability may be an issue because of graphite’s flakiness. Biological pores like AHL or MspA may also be stacked, for example a stack of 10 AHL pores can provide a pore about 60-80 nm thick.

2) *Location of peptidases*. The cleaving action of an enzyme (endopeptidase or exopeptidase) requires it to be in the path of the peptide or fragment emerging from the respective pore (UNP or MNP) on the *trans* side. This can be ensured by covalently attaching the enzyme to the *trans* side of the pore membrane. Such covalent attachment has been discussed for DNA sequencing using two different approaches: exosequencing of mononucleotides^29^ and sequencing by synthesis using heavy tags attached to the bases.^30^ In both approaches an exonuclease or polymerase is attached to the *cis* side of the pore membrane. This could result in significant errors due to cleaved bases or tags being lost to diffusion in the *cis* chamber (deletions) or entering the pore out of order (delete-and-insert).^31^ In the present approach the peptidases are located on the *trans* side so deletions cannot occur. Out-of-order arrivals at the sensing pore (fragments at DNP in *Method 1*; fragments at MNP and residues at DNP in *Method 2*) are precluded as long as the necessary conditions given in Section 3 are satisfied.

3) *Solution pH*. Solution pH plays an important role for two reasons: a) the charge carried by a fragment, which is highly variable and not known in advance, is a function of pH (compare with DNA, where all nucleotide types have approximately the same electron charge of -q with a small variability due to pH); b) its effect on enzyme reaction rates. The choice of pH is a tradeoff between enzyme efficiency and being able to control translocation speeds; this may be determined by experiment.

4) *Fabrication*. A recent review of nanopore sequencing includes notes on fabrication techniques.^27^ Tandem-pore-like structures have been fabricated and used in polymer studies recently. One of these has been used to trap and analyze DNA;^32^ with the *trans*1/*cis*2 chamber functioning like a test-tube. A similar structure has been used to measure the mobility of DNA molecules.^33^ In contrast with conventional nanopore sequencing methods, where the aim is to fabricate thin (∼1 nm) pores that are usually synthetic (for example Si_3_N_4_), as noted earlier the thick (80-100 nm) pores required in the proposed scheme may be more easily fabricated.

6) *Other*. See Appendix.

Successful implementation of the proposed method could lead to a portable easy-to-use low-cost fast-turnaround digital device for protein identification and quantity estimation. Neither analyte labeling nor immobilization is required, only electrical measurements are used, and bandwidths required are within limits. The sample need not be purified to homogeneity; the quantity of a protein in a non-homogeneous mixture can be estimated from the measured number of its identifying peptides. Tandem cells of the required size and shape can be fabricated with available technology, enzymes of the required specificity are available, and sequence data are easily assembled from digital signals output by multiple cells. The assembled partial sequence can be rapidly compared with sequences in a proteome database to find a unique match if one exists. Finally, the low data storage and processing requirements of the proposed method suggest easy integration with hand-held diagnostic and mobile electronic devices (similar to a recently introduced genome sequencer with a USB interface^4^).

## Supplementary information

Data files with identifying sequences for each protein in the human proteome for two sets of cleaving options are available.

## Appendix

### A-1 Translocation statistics of tandem cell

Following [11,12], the mean E(T) and variance σ^2^(T) of the translocation time T over a channel of length L that is reflective at the top and absorptive at the bottom with applied potential difference of V are given by

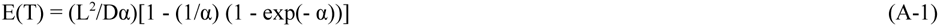

and

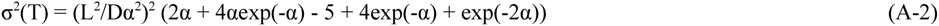

where

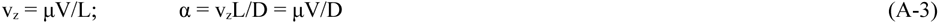

Here v_z_ is the drift velocity due to the electrophoretic force experienced by a charged particle in the z direction, which can be 0, negative, or positive. For v_z_ = 0, these two statistics are

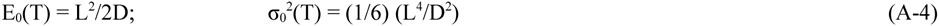

If each section in the double tandem cell is considered independently these formulas can be applied to all the relevant sections: *trans*1/*cis*2 (T = T_*trans*1/*cis*2_; L = L_23_), MNP (T = T_MNP_; L = L_34_), *trans2*/*cis3* (T = T_*trans*2/*cis*3_; L = L_45_), DNP (T = T_DNP_; L = L_56_), and *trans*3 (T = T_*trans*3_; L = L_67_). For an analysis of behavior at the interface between two sections see [11,12].

### A-2 Dependence of particle translocation on solution pH, charge, diffusion constant, and mobility

Equations A-1 through A-4 involve a number of physical-chemical properties of amino acids: electrical charge (itself dependent on solution pH) [34], hydrodynamic radius, diffusion constant, and mobility. The following paragraphs provide a quantitative description of this dependence and allow calculation of fragment properties as they apply to peptide sequencing in a tandem cell with endopeptidase. In particular this information is used in the next section to derive a required condition for effective sequencing.

**Table A-1.**

| Peptide end or amino acid | Amino end | Carboxy end | R | D | C | E | H | K | Y |
| --- | --- | --- | --- | --- | --- | --- | --- | --- | --- |
| **kA value** | 2.34 | 9.69 | 12.48 | 3.86 | 8.33 | 4.25 | 6.0 | 10.53 | 10.07 |

1) The electrical charge carried by a peptide (fragment) P_x_ can be calculated with the Henderson-Hasselbach equation. Let the set of amino acids be **AA** = [A, R, N, D, C, Q, E, G, H, I, L, K, M, F, P, S, T, W, Y, V] where AA[i] is the i-th amino acid, 1 ≤ i ≤ 20. Let the pH value of the solution (electrolyte) be p, kC = kA value of the carboxy end = 9.69, kN = kA value of the amino end = 2.34, N_X_ the number of times residue X occurs in the peptide (X = R, H, K), N_Z_ the number of times residue Z occurs (Z = D, C, E, Y), and kX and kZ the kA values of X and Z respectively. kA values are given by Table A-1. The charge multiplier C_Px_ on the peptide is given by

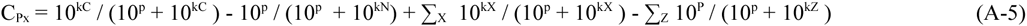

where the summations are over the N_X_ and N_Z_ occurrences of X and Z respectively in P_x_.

2) The hydrodynamic radius R_Px_ of peptide P_x_ = X_1_ X_2_… X_N_ is obtained recursively as follows:

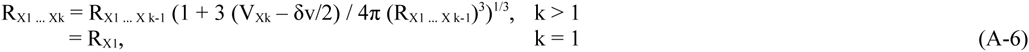

where V_Xk_ and and dv are the van der Waals volumes of X_k_ and a single molecule of water. Hydrodynamic radii of individual amino acids are given in [35] and van der Waals volumes in [36] (both sets of values are reproduced in the Supplement to [12]). This formula holds for small peptides (up to ∼20 residues).

The diffusion constant and mobility of P_x_ are given by

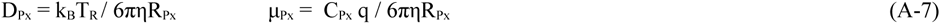

Here k_B_ is the Boltzmann constant (1.3806 × 10^_-23_^ J/K), T_R_ is the room temperature (298° K), η is the solvent viscosity (0.001 Pa.s), q is the electron charge (1.619 × 10^−19^ coulomb), and C_Px_ is a multiplier.

Figure 1 shows the distribution of the number of peptides of length 7 vs mobility μ and μ/D (= a with V set to 1) over all 20^7^ of them.

**Figure A-1.**
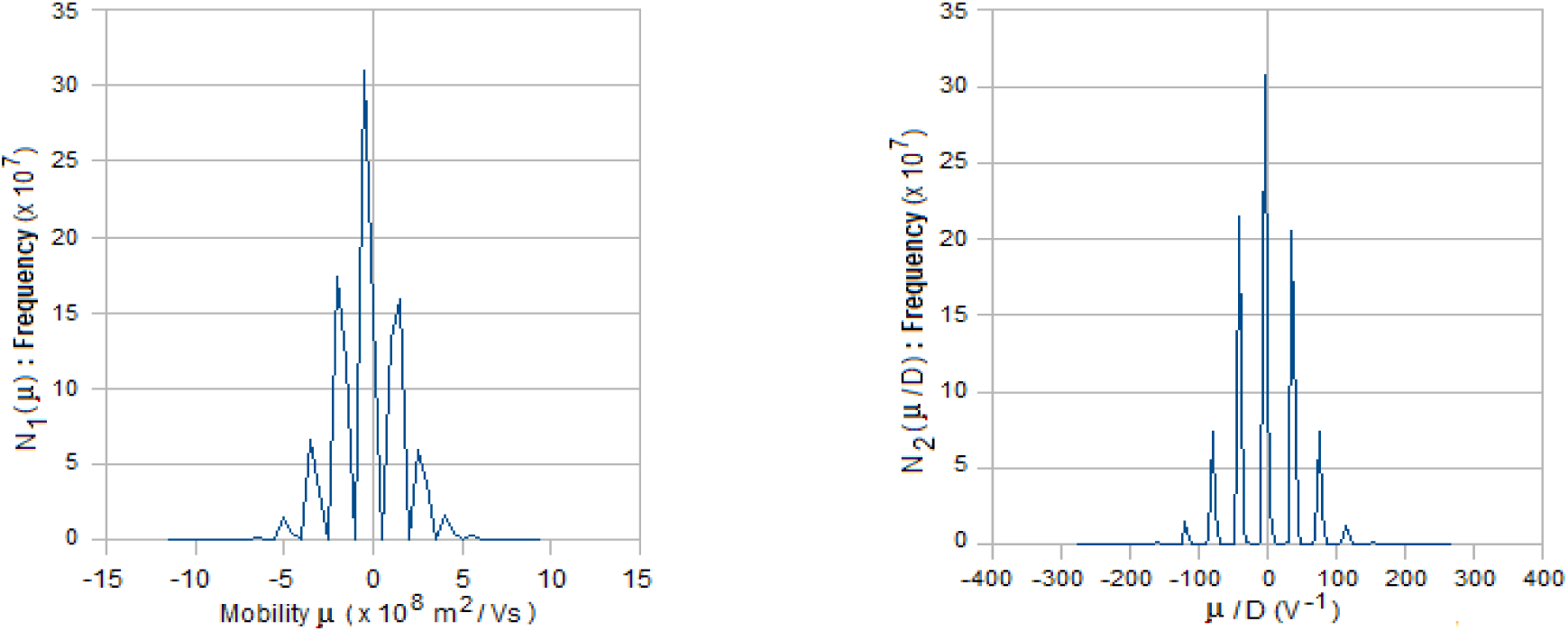
Distribution of number of peptides of length 7 vs mobility μ and the ratio of mobility to diffusion coefficient (μ/D).

### A-3 Calculating the percentage of misses (deletes) due to fast fragments and losses due to slow fragments

For fragments that carry a high negative or positive charge the mean translocation time in Equations A-1 and A-4 can be approximated by

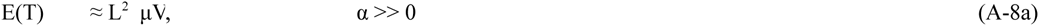

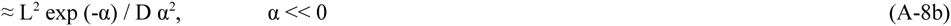

These formulas can be used to estimate the percentage of misses due to fast translocating fragments and slowly moving fragments through DNP in Method 1.

To estimate the former, for a given pore length L, voltage V across the pore, and blockade pulse width approximated by E(T), μ is written as

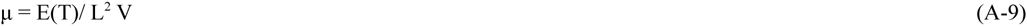

The percentage of misses is given by

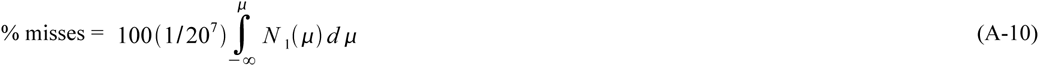

The integral in Equation A-10 is the cumulative frequency for N_1_(μ) corresponding to the μ calculated from Equation A-9. The results are shown in Table A-2 for V = 100 mV, L = 100, 150, 200, and 250 nm, and two pulse widths: E(T) = 7 μs and 10 μs.

To estimate the percentage of losses rewrite Equation A-8b as

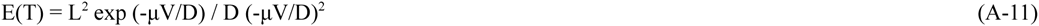

This is an implicit function of two parameters, μ/D and D. To solve for μ/D for a given E(T), L, and V, it is approximated by

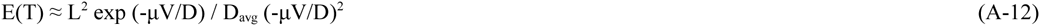

where D_avg_ is the average diffusion coefficient of all 20^7^ peptides of length 7. This is a nonlinear equation in μ/D; the desired root on the real line can be found using standard methods. For a given value of V, the percentage of losses is given by

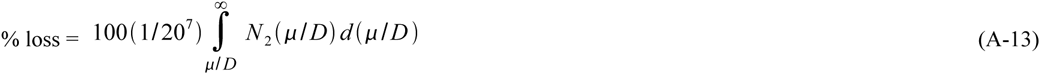

The results are shown in Table A-2 for V = 100 mV and L = 100, 150, 200, and 250 nm, and E(T) = 1 s.

**Table A-2.**

| L (nm) | Pulse width = 7 $\mu\text{s}$ | | | Pulse width = 10 $\mu\text{s}$ | | | Pulse width = 1 s | | |
| --- | --- | --- | --- | --- | --- | --- | --- | --- | --- |
| | $\mu$<br>($\text{m}^2/\text{Vs}$) | $N_1(\mu)$ | Percentage deletions | $\mu$<br>($\text{m}^2/\text{Vs}$) | $N_1(\mu)$ | Percentage deletions | $\mu/D$<br>($\text{V}^{-1}$) | $N_2(\mu/D)$ | Percentage losses |
| 100 | -1.43 | $4.1 \times 10^8$ | 32.09 | -1 | $7.2 \times 10^8$ | 56.27 | $1.6 \times 10^2$ | $2.06 \times 10^5$ | 0.02 |
| 150 | -3.22 | $1.19 \times 10^8$ | 9.3 | -2.25 | $2.93 \times 10^8$ | 22.89 | $1.52 \times 10^2$ | $2.52 \times 10^8$ | 0.19 |
| 200 | -5.72 | $2.97 \times 10^6$ | 0.23 | -4 | $2.2 \times 10^7$ | 1.72 | $1.45 \times 10^2$ | $2.52 \times 10^8$ | 0.19 |
| 250 | -8.94 | $7.4 \times 10^3$ | 0 | -6.25 | $2.5 \times 10^6$ | 0.2 | $1.4 \times 10^2$ | $2.52 \times 10^8$ | 0.19 |

Figures A-2 and A-3 show similar distributions for peptide lengths 12 and 16.

**Figure A-2.**
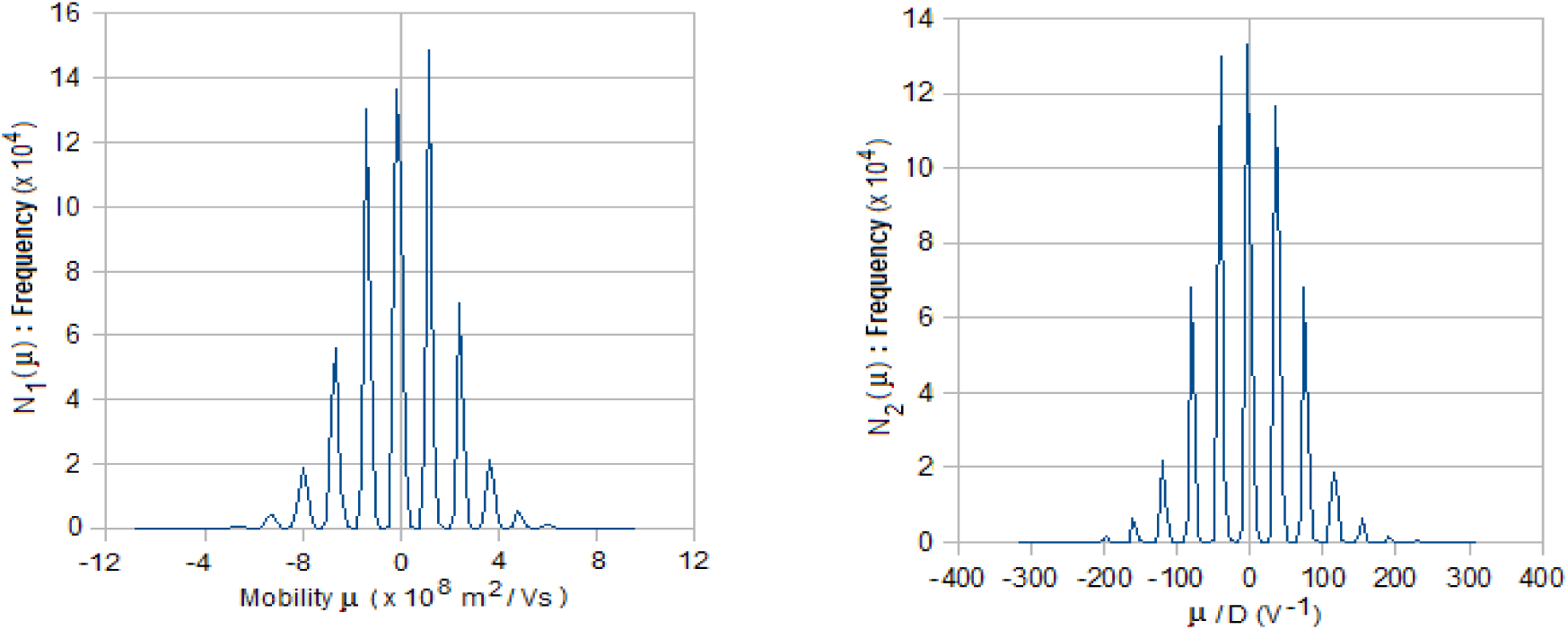
Distribution of number of peptides vs mobility μ and the ratio of mobility to diffusion coefficient (μ/D). Numbers based on 10^6^ randomly generated peptide strings of length 12.

**Figure A-3.**
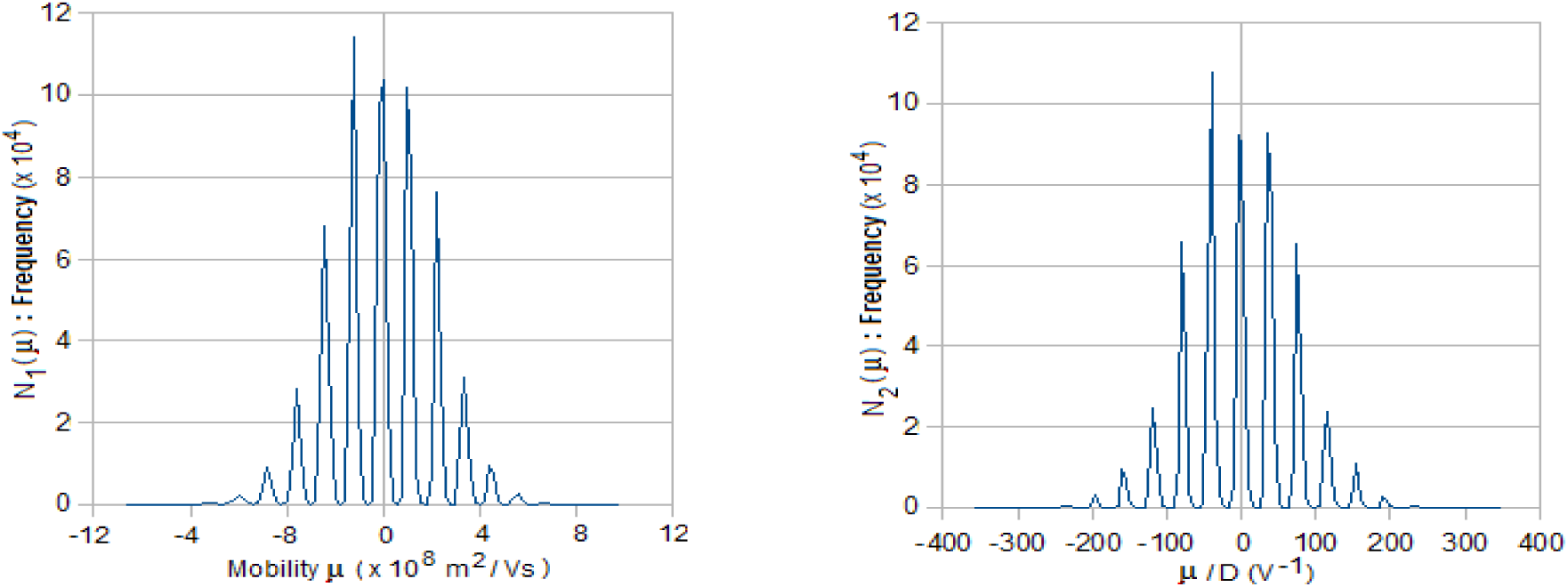
Distribution of number of peptides vs mobility μ and the ratio of mobility to diffusion coefficient (μ/D). Numbers based on 10^6^ randomly generated peptide strings of length 16.

### A-4 Translocation statistics of single residues

The mean and standard deviation of the time taken by a single residue through *trans*2/*cis*3 and DNP (Method 2) are shown in Table A-3 as a function of pH.

**Table A-3.** *Method 2*: Translocation time of single residues through *trans*2/*cis*3 (10^−3^ s) and DNP (10^−6^ s)

| Amino acid | pH=3 |  |  |  | pH=7 |  |  |  | pH=11 |  |  |  |
| --- | --- | --- | --- | --- | --- | --- | --- | --- | --- | --- | --- | --- |
|  | *trans2/cis3* |  | DNP |  | *trans2/cis3* |  | DNP |  | *trans2/cis3* |  | DNP |  |
|  | Mean | Std dev | Mean | Std dev | Mean | Std dev | Mean | Std dev | Mean | Std dev | Mean | Std dev |
| A | 0.1528 | 0.1248 | 5.5557 | 4.8187 | 0.1523 | 0.1244 | 3.8857 | 3.1704 | 0.1501 | 0.1222 | 1.2137 | 0.6792 |
| R | 0.2100 | 0.1721 | 156.44 | 155.54 | 0.2094 | 0.1715 | 79.93 | 78.98 | 0.2062 | 0.1677 | 5.4214 | 4.4501 |
| N | 0.1711 | 0.1398 | 6.2241 | 5.3984 | 0.1707 | 0.1394 | 4.3532 | 3.5518 | 0.1682 | 0.1369 | 1.3597 | 0.7609 |
| D | 0.1731 | 0.1414 | 4.9358 | 4.1143 | 0.1703 | 0.1386 | 1.3264 | 0.7297 | 0.1678 | 0.1362 | 0.7536 | 0.3153 |
| C | 0.1643 | 0.1342 | 5.9734 | 5.1809 | 0.1637 | 0.1336 | 3.8589 | 3.0969 | 0.1589 | 0.1290 | 0.7143 | 0.2990 |
| Q | 0.1855 | 0.1516 | 6.7462 | 5.8513 | 0.1850 | 0.1510 | 4.7184 | 3.8497 | 0.1823 | 0.1484 | 1.4738 | 0.8248 |
| E | 0.1802 | 0.1472 | 5.8700 | 5.0063 | 0.1771 | 0.1441 | 1.3803 | 0.7596 | 0.1745 | 0.1416 | 0.7835 | 0.3278 |
| G | 0.1332 | 0.1089 | 4.8456 | 4.2028 | 0.1329 | 0.1085 | 3.3891 | 2.7651 | 0.1309 | 0.1066 | 1.0586 | 0.5924 |
| H | 0.2036 | 0.1668 | 151.08 | 150.21 | 0.2002 | 0.1635 | 6.0541 | 5.0998 | 0.1969 | 0.1603 | 1.5924 | 0.8912 |
| I | 0.1861 | 0.1520 | 6.7671 | 5.8694 | 0.1856 | 0.1515 | 4.7330 | 3.8617 | 0.1828 | 0.1488 | 1.4783 | 0.8273 |
| L | 0.1947 | 0.1591 | 7.0804 | 6.1411 | 0.1942 | 0.1585 | 4.9521 | 4.0404 | 0.1913 | 0.1557 | 1.5468 | 0.8656 |
| K | 0.2153 | 0.1764 | 160.35 | 159.42 | 0.2147 | 0.1758 | 81.84 | 80.87 | 0.2090 | 0.1703 | 2.1084 | 1.2987 |
| M | 0.1769 | 0.1445 | 6.4329 | 5.5795 | 0.1764 | 0.1440 | 4.4993 | 3.6710 | 0.1738 | 0.1415 | 1.4053 | 0.7865 |
| F | 0.1924 | 0.1572 | 6.9969 | 6.0686 | 0.1919 | 0.1567 | 4.8937 | 3.9928 | 0.1890 | 0.1539 | 1.5285 | 0.8554 |
| P | 0.1539 | 0.1257 | 5.5975 | 4.8549 | 0.1535 | 0.1253 | 3.9150 | 3.1942 | 0.1512 | 0.1231 | 1.2228 | 0.6843 |
| S | 0.1585 | 0.1295 | 5.7646 | 4.9998 | 0.1581 | 0.1291 | 4.0318 | 3.2896 | 0.1557 | 0.1268 | 1.2593 | 0.7048 |
| T | 0.1746 | 0.1426 | 6.3494 | 5.5071 | 0.1741 | 0.1422 | 4.4409 | 3.6233 | 0.1716 | 0.1397 | 1.3871 | 0.7763 |
| W | 0.2010 | 0.1642 | 7.3102 | 6.3404 | 0.2005 | 0.1637 | 5.1128 | 4.1715 | 0.1975 | 0.1608 | 1.5970 | 0.8937 |
| Y | 0.2050 | 0.1675 | 7.4564 | 6.4672 | 0.2045 | 0.1669 | 5.2070 | 4.2471 | 0.1987 | 0.1613 | 0.9359 | 0.4014 |
| V | 0.1907 | 0.1558 | 6.9342 | 6.0143 | 0.1901 | 0.1553 | 4.8499 | 3.9570 | 0.1874 | 0.1525 | 1.5148 | 0.8477 |

*trans2/cis3*: height = 0.5 $\mu\text{m}$ radius = 0.5 $\mu\text{m}$ $V_{23} = 1.2$ mV DNP: height = 80 nm radius = 2 nm $V_{34} = 140$ mV

### A-5 Derivation of necessary conditions for effective sequencing

It is now shown that the necessary conditions applicable to each of the two methods are satisfied by a large majority (∼80% in most cases) of peptide sequences of a given length for a set of typical parameter values. In the following translocation time distributions are assumed to have 6σ support (σ = standard deviation). The following parameter values are assumed: V_05_ = ∼115 mV (*Method 1*); V_07_ = ∼180 mV (*Method 2*); detector resolution = 1 μs; pore (DNP, MNP) conductance = ∼1 nS; pH = 7.0; *trans*1/*cis*2 height = *trans*2/*cis*3 height = 0.5 μm, UNP length = MNP length = 10 nm. V_05_ divides as V_01_ = V_23_ = V_45_ ≈ 1.6 mV, V_12_ ≈ 10 mV, and V_34_ ≈ 100 mV. V_07_ divides as V_01_ = V_23_ = V_45_ = V_67_ ≈ 1.5 mV, V_12_ = V_34_ ≈ 15 mV, and V_56_ = 140 mV. Let T_exo-min_, T_endo-min-2_, and T_endo-min-1_ be the minimum reaction time intervals for the exopeptidase in *Method 2*, the endopeptidase in *Method 2*, and the endopeptidase in *Method 1* respectively.

**Method 2**. Referring to Section 3 in the main text *Conditions 1a*, *1b*, *2a*, *2b*, and *3* have to be satisfied. From Table A-3, with pH = 7.0, DNP height = 80 nm, and V_56_ = 140 mV, the fastest amino acid is Asp (D) with a translocation time of ∼1.33 μs > 1 μs. This satisfies *Condition 3*.

Let X_1_ and X_2_ be two residues cleaved in succession by the exopeptidase. *Conditions 1a*, *1b*, and *2a* are satisfied if

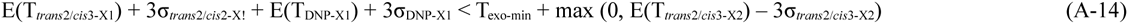

From columns 6 and 7 in the same table the second term in the inequality on the right is 0, leading to

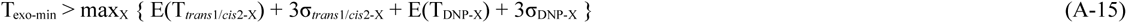

over all X. The maximum occurs for X = K (Lys), with E(T_*trans*2/*cis*3-X_) = 0.21×10^−3^, σ_*trans2/cis3-X*_= 0.18×10^−3^, E(T_DNP-X_) = 82×10^−6^, and σ_DNP-X_ = 81×10^−6^, leading to

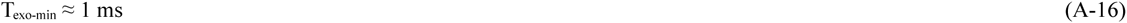

More generally the rate can be calculated for each residue type in a similar way. More generally Figure A-4 shows the mean blockade pulse widths due to single residues in DNP for all 20 residue types for three different lengths of DNP, while Figure A-5 shows T_exo-min_ vs residue type for DNP length = 80 nm.

A peptide that has threaded through UNP encounters the endopeptidase in *trans*1/*cis*2 and is cleaved into fragments. The latter translocate through *trans*1/*cis*2 and thread through MNP to be cleaved by the exopeptidase on the downstream side. Consider two successive fragments F_1_ and F_2_. Let L_F1_ be the length of a fragment F_1_. The delay due to cleaving of F_1_ into single residues by the exopeptidase is L_f1_T_exo-min-2_. *Conditions 1a* and *2b* will be satisfied if

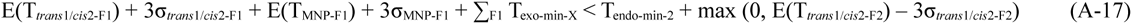

where T_exo-min-X_ is the cleavage time for residue X and the summation is over all L_F1_ residues in F_1_. In the second term on the right side of the inequality, σ_*trans*1/*cis*2-F2_ ≈ E(T_*trans*1/*cis*2-F2_), so that max (0, E(T_*trans*1/*cis*2-F2_) – 3σ_*trans*1/*cis*2-F2_) = 0; this leads to

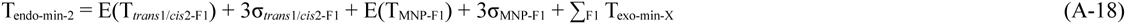

Figure A-6 shows the distribution of T_endo-min-2_ with 10^6^ random peptide sequences with residues in a sequence drawn from a uniform distribution for three different fragment lengths.

**Method 1**. The development is similar to that for *Method 2*. Thus *Conditions 1a*, *2a*, and *3* have to be satisfied. With two successive fragments F_1_ and F_2_ cleaved by the endopeptidase, *Conditions 1a* and *2a* require

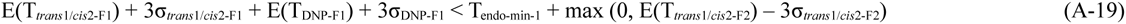

As before the second term on the right is 0 because σ_*trans*1/*cis*2-F2_ ≈ E(T_*trans*1/*cis*2-F2_) which leads to

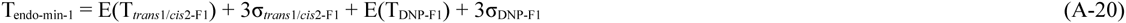

Figure A-7 shows the distribution of fragment pulse widths for three different fragment lengths. Figure A-8 shows T_endo-min-1_ for 10^6^ random samples of length 12. For the vast majority of sequences T_endo-min-1_ is < 3 ms. The curve is to the left of the corresponding curve in *Method 2* (Figure A-6, red) because the endopeptidase reaction times in the latter include the delay due to the cleaving of residues in a fragment by the exopeptidase (although this is not strictly necessary because the pulses are only counted so they can arrive in any order). The distribution of pulse widths > 12 μs due to fragments of length = 12 vs the endopeptidase reaction time is shown in Figure A-9. For nearly 80% of the sequences (with blockade pulses in which LF transitions can be counted) T_endo-min-1_ < 1 ms. In comparison the percentage of pulses that may not be counted correctly is relatively small at ∼17%. Incidentally the curves in Figures A-6 are similar in shape and range to reaction rate graphs for the enzyme Exonuclease I [37].

**Figure A-4.**
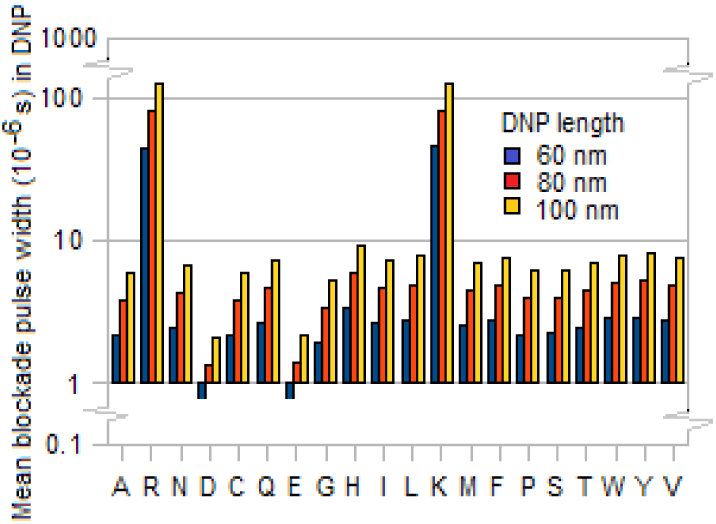
Mean blockade pulse width (μs) in DNP of different lengths for the 20 individual amino acids in *Method 2*. *trans2* height = 0.5 μm, V_56_ = 140 mV, V_45_ = 1.2 mV, pH = 7.

**Figure A-5.**
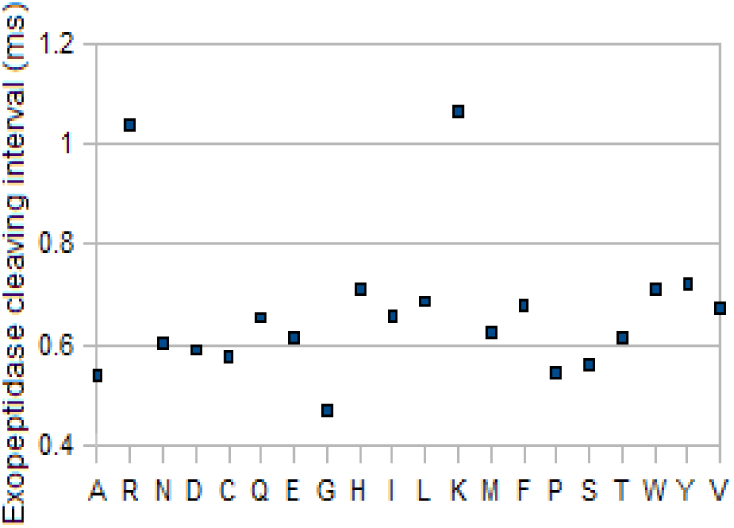
Distribution of T_exo-min_ (ms) for each amino acid type in *Method 2*. DNP height = 80 nm, *trans*2 height = 0.5 μm, V_56_ = 140 mV, V_45_ = 1.2 mV, pH = 7.

**Figure A-6.**
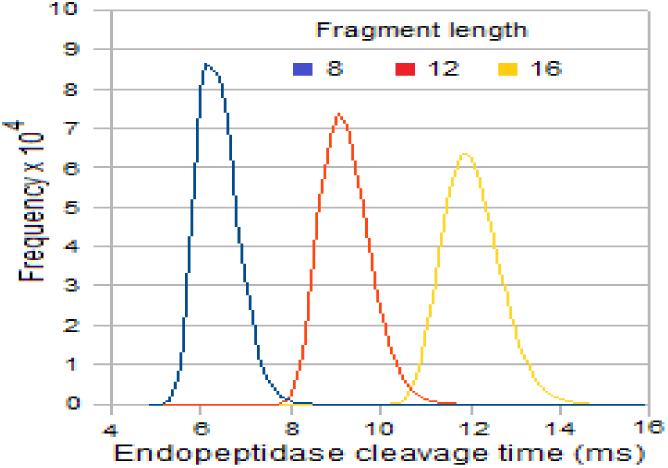
Frequency distribution of T_endo-min-2_ (ms) for different fragment lengths L_F_ in *Method 2*. MNP height = 10 nm, trans1 height = 0.5 μm, V_23_ = 1.2 mV, V_34_ = 20 mV, pH = 7.

**Figure A-7.**
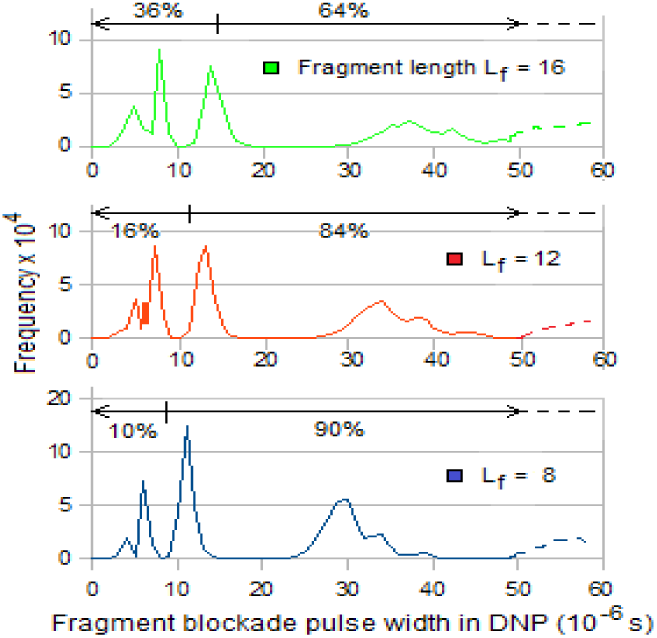
Frequency distribution of fragment pulse widths (μs) for different fragment lengths in *Method 1*. DNP height = 150 nm, *trans*1 height = 0.5 μm, V_34_ = 100 mV, V_23_ = 1.2 mV, pH = 7.

**Figure A-8.**
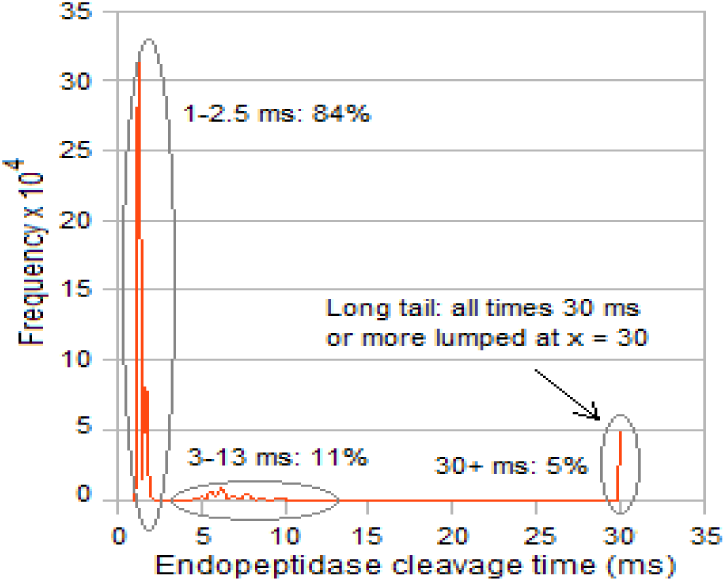
Frequency distribution of T_endo-min-1_ (ms) for fragment length = 12 in *Method 1*. DNP height = 150 nm, *trans*1 height = 0.5 μm, V_34_ = 100 mV, V_23_ = 1.2 mV, pH = 7.

**Figure A-9.**
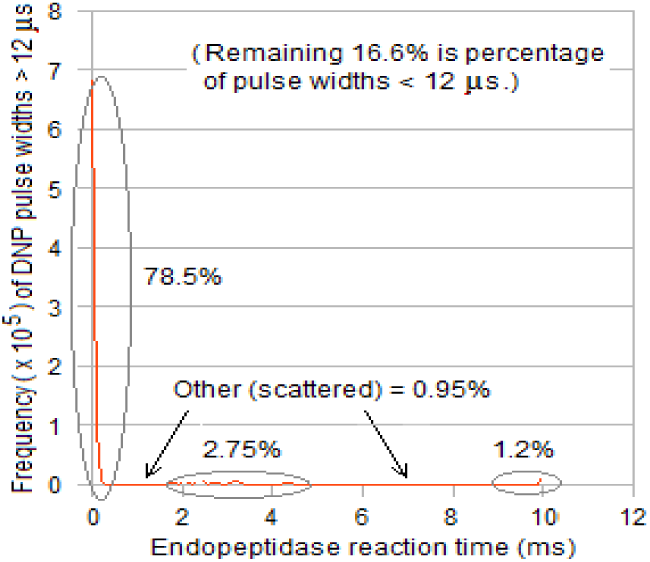
Distribution of pulse widths > 12 μs for fragment length = 12 vs T_endo-min-1_ (ms) in *Method 1*. DNP height = 150 nm, *trans*1 height = 0.5 μm, V_34_ = 100 mV, V_23_ = 1.2 mV, pH = 7.

### A-6 Peptidases and chemicals for cleaving and their specificities

Table A-4 is a summary of selected chemicals and peptidases for use in cleaving of the unknown protein or peptides generated from it at desired locations; it is adapted from [14]. The following notation is used for cleavage sites on a substrate [38]:

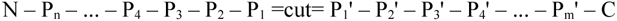

where – represents a peptide bond, N is the N-terminal end, and C is the C-terminal end.

**Table A-4.** *Selected chemical agents for cleaving a protein after or before a specific amino acid (Stage 1)*

| Chemical | Cleavage point and target | Chemical | Cleavage point and target(s) |
| --- | --- | --- | --- |
| BNPS-Skatole | $N - \dots - \text{Trp} = \text{cut} = \dots - C$ | Hydroxylamine | $N - \dots - \text{Asn} = \text{cut} = \text{Gly} - \dots - C$ |
| Cyanogen bromide (CNBr) | $N - \dots - \text{Met} = \text{cut} = \dots - C$ | Iodosobenzoic acid | $N - \dots - \text{Trp} = \text{cut} = \dots - C$ |
| Formic acid | $N - \dots - \text{Asp} = \text{cut} = \dots - C$ | NTCB + Ni | $N - \dots = \text{cut} = \text{Cys} - \dots - C$ |

Selected peptidases for cleaving a peptide after or before one or more specific amino acids (Stage 1 or Stage 2) (Alternative target, usually less probable, is in parentheses.)
| Peptidase | Cleavage point | Target X (or Z) | Peptidase | Cleavage point | Target X |
| --- | --- | --- | --- | --- | --- |
| ArgC proteinase | $N - \dots - X = \text{cut} = \dots - C$ | X = Arg | ArgC endopeptidase | $N - \dots - X = \text{cut} = \dots - C$ | X = Arg (Lys) |
| AspN endopeptidase | $N - \dots = \text{cut} = X - \dots - C$ | X = Asp (Glu) | LysC endopeptidase | $N - \dots - X = \text{cut} = \dots - C$ | X = Lys (Asn) |
| Trypsin | $N - \dots X = \text{cut} = Z - \dots - C$ | X = Arg or Lys Z ≠ Pro | LysC lysyl endopeptidase | $N - \dots - X = \text{cut} = \dots - C$ | X = Lys |
| LysN peptidyl metalloendopeptidase | $N - \dots = \text{cut} = X - \dots - C$ | X = Lys | Neutrophil elastase | $N - \dots - X = \text{cut} = \dots - C$ | X = Val or Ala |
| Glutamyl endopeptidase | $N - \dots - X = \text{cut} = \dots - C$ | X = Glu | | | |

### A-7 Additional notes and references

1) Order *of fragment entry into DNP*. A fragment can enter DNP amino-end first or carboxy-end first. However the order is not important as the information sought is the number of residues, not their identity or sequence.

2) *Order of entry of peptide into UNP*. The assembly algorithm described in Section 2 implicitly assumes that entry of a peptide into UNP in each of the cells is all of them either N-terminal first or C-terminal first. This is a reasonable assumption because of the charged X-header. However, there is a non-zero probability that the peptide may enter wrong end first, so some of the fragment length lists obtained will be in the reverse order. The assembly algorithm needs to be modified to take this into account.

3) *Applied voltage and current levels*. Blockades are of ionic current flow through the pore due to K+ and Cl-ions in the electrolyte; with V = ∼100 mV this current is ∼100 pA (≈ G_pore_V, where G_pore_ is the conductance of the pore, typically 1 nS for a pore ∼10 nm long), usually adequate for measuring blockades [6]. With longer pores blockade levels may be lower. In the presence of noise there is a tradeoff between detectable pulse amplitude changes and translocation speed. While a higher voltage results in a higher blockade current and higher signal-to-noise ratios (SNR), it also causes a fragment or residue with high negative charge to translocate through DNP at a rate that exceeds 1/T_detector_, and one with high positive charge to translocate too slowly, resulting in misses or ‘loss’ to diffusion respectively. These extremes have been estimated in Section A-3 above. (The upper limit to the applied voltage is set by the breakdown field for the electrolyte, typically ∼70 MV/m.)

4) *Entropy barriers*. It is assumed that the entropy barrier [6] faced by a fragment during its entry into DNP (*Method 1*) or MNP (*Method 2*) is negligible, in part because short peptides have been considered. Long peptides may form secondary structures and also ball up, impeding entry into a pore. In this case, the barrier may not be negligible; it can be taken into account by increasing the minimum cleaving intervals required of the enzymes. The taper in *trans*1/*cis*2 and *trans*2/*cis*3 (Figures 1 and 2) also helps lower the entropy barrier. Based on the computational results discussed above, the two methods presented here appear well suited to sequencing of peptides with 12-16 residues. (Compare with the optimum peptide length of ∼20 in an efficient mass spectrometer [3].)

5) *Independence of cells*. Each cell targets a different amino acid and operates independent of the other cells. This means that the cell can be independently optimized for enzyme reaction rates, applied voltage, pH value, etc.

6) *Sticky fragments/residues*. The problem of fragments or residues sticking to pore or compartment walls may be resolved through the use of non-stick additives [39] or wall coatings [40].

7) *Sequencing with the potential reversed*. A peptide can be sequenced with the applied potential reversed, which speeds up fragments with positive charge and slows down those with negative charge; neutral fragments are not affected. (If the pore is ion-sensitive, one with the appropriate sense may be used.) Merging the two sets of data can lead to improvements in detection and correction of errors, but this is only for charged fragments. The error can be minimized over all fragments, charged or neutral, by experimentally varying the pH and finding the pH value that yields the best results.

8) *Hafnium oxide pores*. Recent studies using high bandwidth (∼4 MHz) detectors have shown that a HfO_2_ membrane < 10 nm thick can slow down translocating DNA molecules [41]. (The slowdown is believed to be due to interactions of the DNA with the walls of the pore.) At the present time, however, fabrication seems to require an inordinate amount of time.

For other implementation-related issues affecting tandem cells see discussions in [11,12].

